# Rarity of microbial species: In search of reliable associations

**DOI:** 10.1101/358077

**Authors:** Arnaud Cougoul, Xavier Bailly, Gwenaëel Vourc’h, Patrick Gasqui

## Abstract

The role of microbial interactions on the properties of microbiota is a topic of key interest in microbial ecology. Microbiota contain hundreds to thousands of operational taxonomic units (OTUs), most of which are rare. This feature of community structure can lead to methodological difficulties: simulations have shown that methods for detecting pairwise associations between OTUs (which presumably reflect interactions) yield problematic results. The performance of association detection tools is impaired for a high proportion of zeros in OTU table. Here, we explored the statistical testability of such associations given occurrence and read abundance data. The goal was to understand the impact of OTU rarity on the testability of correlation coefficients. We found that a large proportion of pairwise associations, especially negative associations, cannot be reliably tested. This constraint could hamper the identification of candidate biological agents that could be used to control rare pathogens. Consequently, identifying testable associations could serve as an objective method for trimming datasets (in lieu of current empirical approaches). This trimming strategy could significantly reduce the computation time and improve inference of association networks. When OTU prevalence is low, association measures for occurrence and read abundance data are correlated, raising questions about the information actually being captured.

## Introduction

Microbiota play key roles in ecosystem processes, from eukaryote physiology [1] to global biogeochemical cycles [2]. Research often focuses on comparing microbiota found in similar environments to identify the major forces shaping their structure [3] and function [4]. Microbial interactions are probably one such force [5, 6].

The most common technique for describing microbiota is 16S rRNA sequencing [7]. Association network analysis is then often employed to characterize potential microbial interactions [8]. Association networks require identifying pairwise associations between the occurrence or abundance of bacterial operational taxonomic units (OTUs) [9].

However, microbiota frequently contain hundreds to thousands of OTUs, most of which are rare [10, 11]. A typical matrix describing the abundance of OTUs among similar microbiota thus includes a high proportion of zeros. Simulations illustrated that an excess of zero impairs the efficiency of association network analysis [12, 13]. To avoid this, rare OTUs are filtered out before such analysis. Current trimming procedures are empirical in nature and restrictive. They may rely on OTU prevalence [12, 14], mean abundance [15], or diversity [16]. Moreover, simulations suggested that association network analyses more efficiently detect negative relationships (amensal, competitive) than positive ones (i.e., mutual, commensal) [12]. It is not clear yet wether this is due to the distribution of OTU prevalence.

Precise definition of the conditions under which positive and negative associations are reliably tested can help improve current studies on microbial interactions. This can help define a study plan that will provide sufficient statistical power. This can evidence potential avenues for improving data analysis. This can help interpret association network analysis, depending on the limitations of these approaches.

Below, we analyzed the effect of low OTU prevalences observed in microbiota on associations measures based on occurrence data and read abundance data. More specifically, we computed the extrema of correlation coefficients depending on prevalences for both data types. These extrema were used to define which associations between OTUs could be reliably tested. We finally compared the results obtained from both data types. This allowed (i) to define to what extent prevalence and sample size affect the association studies in microbiota, (ii) to demonstrate that negative interactions can not be captured in most cases, (iii) to show that the added value obtained from analyzing abundance data compared to occurrence data is limited. Results are discussed in the light of current analysis procedures and tools to identify potential solutions to the issues we evidenced.

## Materials and methods

The distribution of association statistics are affected by prevalence. This lead to problems to test correlation coefficients. For instance, the statistic’s minimum and/or maximum can fall within the expected confidence interval obtained from the classical distributions used to approximate expected values. This issue can arise with both occurrence and abundance data.

### Model for occurrence data

First, we explored how to define testability when occurrence data are used. Regarding tests based on the expected distribution of an association statistic, we employed the Phi coefficient *ϕ* [17], a measure of association between two binary variables *X*_*A*_ and *X*_*B*_:

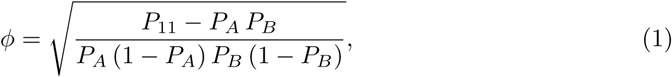

where *P*_*A*_, *P*_*B*_ are the prevalence values for two OTUs, *X*_*A*_ and *X*_*B*_, and *P*_11_ is the prevalence of their co-occurrence. The extrema of Phi [18] depend exclusively on *P*_*A*_ and *P*_*B*_ (S1 Fig).

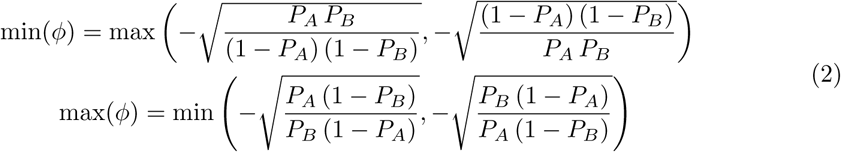

Under the null hypothesis (*H*_0_) that the occurrences of *X*_*A*_ and *X*_*B*_ are independent, Phi can be approached thanks to the Pearson’s chi-squared test:

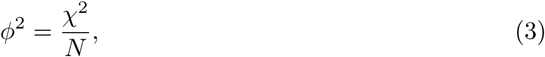

where *N* is the total number of samples and *χ*^2^ is a chi-squared distribution with 1 degree of freedom [19]. This last distribution is thus used to build a confidence interval to test departure from the independence hypothesis. Furthermore, we can describe cases where associations can not be tested reliably based on this confidence interval, because the genuine minimum and/or maximum of *ϕ* fall within this confidence interval.

For occurrence data, limitations on testability can also be studied with exact tests based on the possible combinations of associations (like the Fisher’s exact test, see Part 2.7 in S1 Appendix). For fixed prevalences, the probability of observing the minimum or maximum number of co-occurrences may be higher than the alpha level (canonically at 5%) [20, 21]. In such a case, a negative or positive association, respectively, can not be significantly detected.

### Model for read abundance data

Second, we explored how to define testability when read abundance data are used. We employed the Pearson correlation coefficient [22], a measure of association between two continuous variables, *X*_*A*_ and *X*_*B*_.

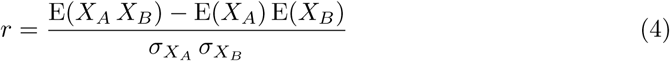

We demonstrated that the minimum of Pearson correlation coefficient depends only on OTU prevalence (see proof in Part 3 in S1 Appendix and illustration S2 Fig).

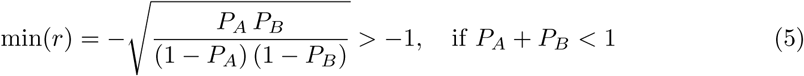

We can then set up a confidence interval from the following assumption. When *X*_*A*_ and *X*_*B*_ follow two uncorrelated normal distributions,

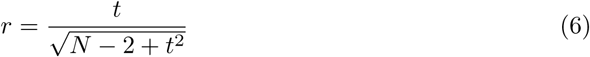

where *t* has a Student’s *t*-distribution with degrees of freedom *N* − 2.

We provide a detailed description of how prevalence affects the testability of associations for both data types in a detailed supplementary material (S1 Appendix).

To estimate the proportion of unreliable tests, we considered two distributions for the OTU prevalence: (i) an uniform law to study the influence of sample size *N* and prevalence *P*_*A*_, *P*_*B*_; (ii) a truncated power law to take into account the real structure of microbiota data. We also compared the results on the testability limits for the two types of data and highlighted a correlation between the two associated measures.

## Results

### Testability given a uniform distribution of prevalence

When occurrence data were used, four inequations (Eqs (7ߝ10) in S1 Appendix) defined reliable tests based on the chi-squared distribution depending on OTU prevalences (Fig 1A). The proportion of non-testable associations (i.e., neither positive nor negative correlations could ever be significant) rapidly fell as *N* increased (Fig 1B). The proportion of associations with partial testability (i.e., either only positive or negative correlations could ever be significant) never exceeded 0.25 (Fig 1B). When *N* = 300, the proportion of fully testable associations (both positive and negative correlations could be significant) exceeded 0.80 (Fig 1B). We showed by simulations that the proportion of Fisher’s exact test affected by prevalence are similar compared to the analytical results presented above (S3 Fig).

**Fig 1.**
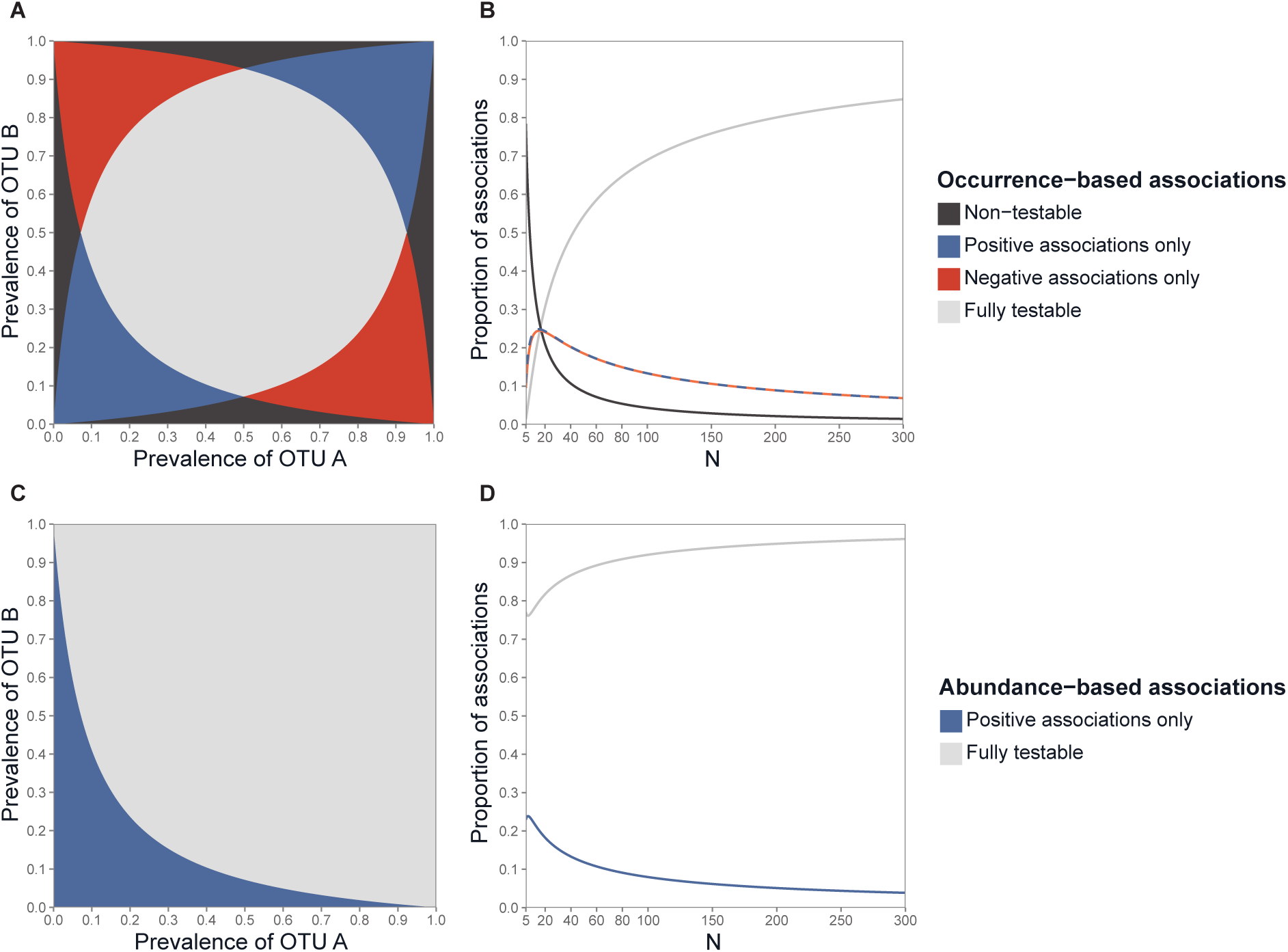
Testability of pairwise associations for the occurrence and for the read abundance data. For the occurrence data: testability zones defined by OTU prevalence (**A**), and the proportion of testable associations as a function of N assuming prevalence follows a uniform distribution (**B**). For the read abundance data: testability zones defined by OTU prevalence (**C**), and the proportion of testable associations as a function of *N* assuming prevalence follows a uniform distribution (**D**). The alpha level for the tests is 5%. For (A) and (C), *N* = 50.

When read abundance data were used, some negative correlations were not testable based on the Student’s distribution (Eq (33) in S1 Appendix and Fig 1C). This problem became less pronounced as *N* increased, and the proportion of testable associations reached 0.95 at *N* = 300 (Fig 1D).

### Testability given observed community structure

Prevalence distributions are highly unbalanced in microbiota because of the large number of rare OTUs (Fig 2A). Accordingly, we modelled OTU prevalence using a truncated power law distribution; the latter reflects observed community structure (Part 5 in S1 Appendix and S4 Fig). OTU prevalence was fitted according to a truncated power law, with k ranging from *−*2 to *−*1: the smaller *k*, the higher the proportion of rare species. Such distribution imply that for most pairs of OTUs, the two OTUs have a low prevalence (Fig 2B).

**Fig 2.**
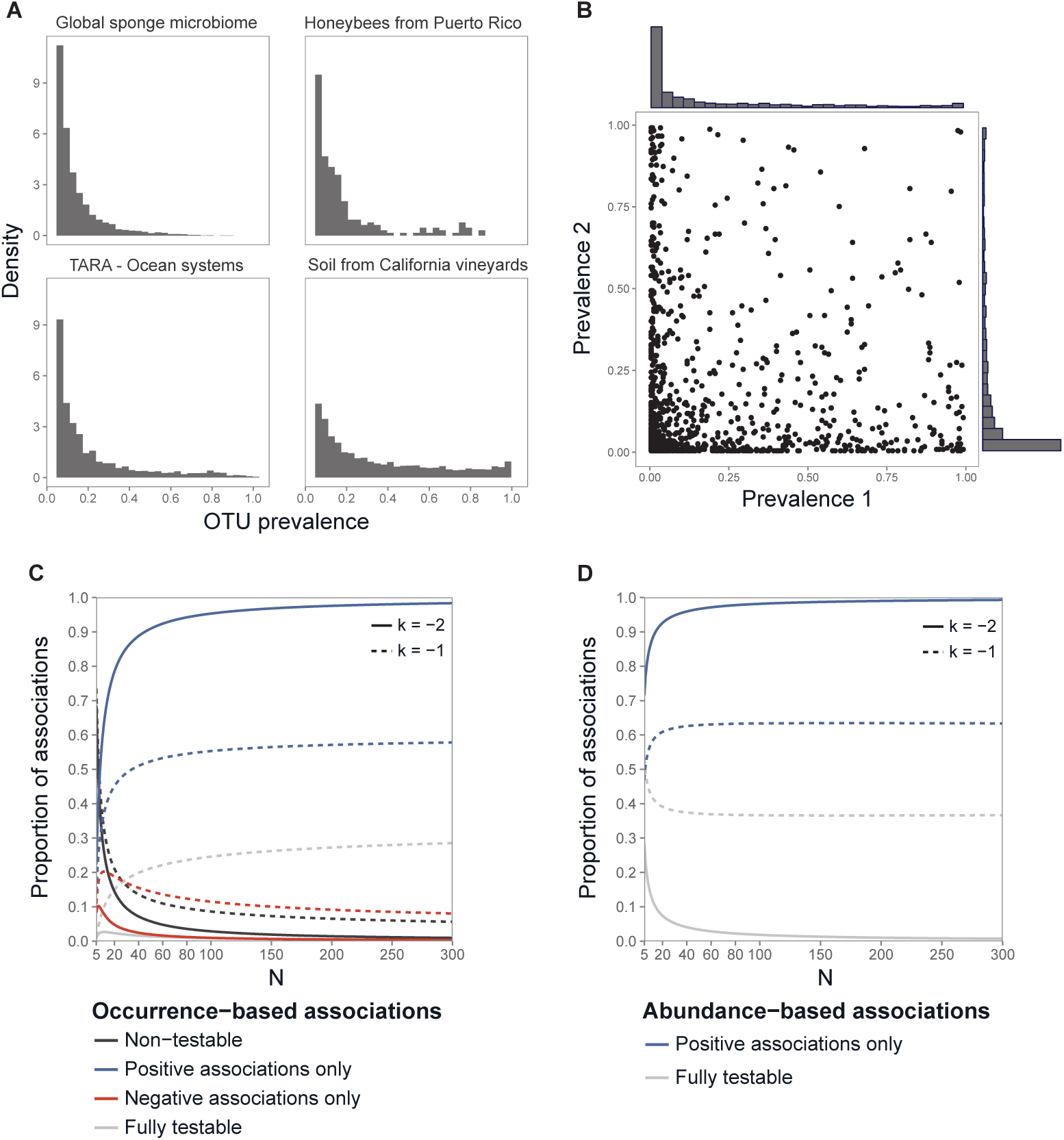
Association testability under realistic conditions of microbial community structure. (**A**) Histograms of OTU prevalence in several microbiota characterized by 16S rRNA sequencing. Data were taken from the QIITA database and the TARA Ocean Project. The microbiota are described elsewhere (Part 5 in S1 Appendix). To better illustrate the skewed distributions, only prevalence values of greater than 5% were included. (**B**) Distribution of all OTU prevalence pairs from microbiota data of the soil from California vineyards. Each point represent a pair of OTU prevalence. For the occurrence data (**C**) and the read abundance data (**D**): proportion of testable associations as a function of *N* when *k* = *−*2 or *−*1.

For the occurrence data, there was thus a large proportion of associations for which negative correlations could never be significant (*>* 0.50 for *k* = −1, *>* 0.90 for *k* = −2); this proportion increased as *N* increased (Fig 2C). This counter-intuitive result is due to the accumulation of rare OTUs as N increases under the power law assumption. Fewer than 10% of associations were non-testable when *N* was greater than 50 (Fig 2D).

For the read abundance data, when *N* = 100, a large and extremely large proportion of negative correlations were untestable when *k* = −1 (proportion: 0.60) and *k* = −2 (proportion: 0.95), respectively (Figs 2D).

### Comparison between the two data types

We finally compared the association statistics for both data types under conditions of low OTU prevalence such as those observed in the actual microbiota data (Part 4 in S1 Appendix). A formal decomposition of variance and covariance illustrates the structural relationship of the correlation coefficients calculated from the occurrence and read abundance data (Eq (2), Part 1 in S1 Appendix). The observed values of the Phi coefficient *ϕ* and the Pearson correlation *r* among OTUs pairs of microbial datasets, (Fig 3A) illustrated that the minimum of the statistics is particularly affected as explained above. Furthermore a correlation is observed between the two measures on real microbial datasets (cor = 0.78 and *R*^2^ = 0.62 on honeybees microbiota data, Fig 3A). Simulations allowed us to investigate more precisely the expected correlation between the two measures. The association tests that can be performed using occurrence versus read abundance data tend to be similar, and prevalence influences association testability in the same way. More specifically, association measures for the two data types become correlated as prevalence decreases (Fig 3B).

**Fig 3.**
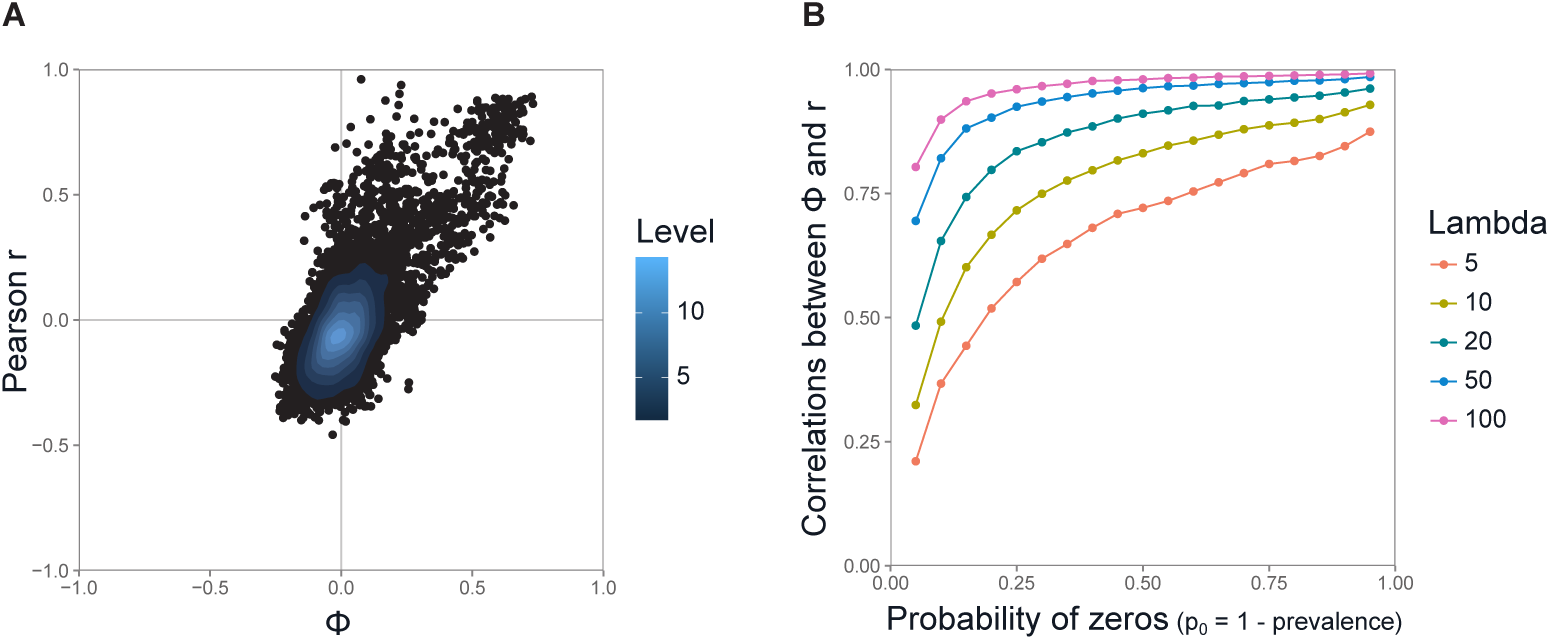
Correlation between the Phi coefficient and the Pearson coefficient. (**A**) Correlation from honeybees microbiota data (Part 5 in S1 Appendix). Each point corresponds to the association coefficients of a pair of OTUs. (**B**) Correlation computed from simulations of OTU abundances modelled by a zero-inflated Poisson (ZIP) distribution (Part 4.2 in S1 Appendix). Parameters are the probability of structural zeroes, *p*_0_, and the value of the Poisson parameter, *λ*. In biological terms, the probability of structural zeroes would correspond to the prevalence (prevalence= 1 *− p*_0_), and the Poisson parameter would correspond to read abundance. For each pair of *p*_0_ and *λ* values, we generated 100 samples of two hypothetical OTUs, *X*_*A*_ and *X*_*B*_, whose abundances followed a zero-inflated Poisson distribution with those parameter values. We then calculated *ϕ* and *r* for these samples. The correlation between the two coefficients was assessed by repeating this process 10^5^ times.

## Discussion

We showed that it is impossible to reliably test a large proportion of the pairwise associations between OTUs in microbiota. Indeed, assuming realistic community structure (i.e., most OTUs are rare), negative correlations could not be tested for most associations. This finding clarifies previous modelling results [12] and underscores a major analytical challenge in this domain. From a practical perspective, for example, this constraint could hamper the identification of candidate biological agents that could be used to control rare pathogens.

This result is amplified by the dependencies within the correlation matrix. There are naturally more positive coefficients than negative coefficients in the correlation matrix. If A and B as well as B and C are negatively correlated, A and C will be more likely to be positively correlated. It is not clear that partial correlations could cope with this issue, especially since a partial negative correlation would be difficult to estimate.

Applying more stringent standards (i.e., analysing only fully testable associations) could drastically reduce the number of tests required to infer an association network. Associations with partial testability (i.e., positive or negative association cannot be tested) could be included, but significance thresholds would need to be adjusted to avoid spurious correlations. We thus propose that identifying testable associations could serve as an alternative to current, empirical strategies for trimming microbiota datasets. By limiting test number, this approach could help control the overall risk of statistical errors and speed up computation time. It could be implemented by forcibly introducing zeros in the correlation matrix during the inference of an association network.

We found that association testability tends to be similar for occurrence and read abundance data. More specifically, association measures calculated using the two data types become correlated as prevalence decreases. This raises questions about the information actually being captured by current methods for quantifying OTU associations. These questions have both computational implications, questioning the ability of current models to make the most of abundance data, and biological implications, because different biological processes of interactions could be revealed by the two data types. Fitting abundances to zero-inflated distributions [23, 24], that aim at decoupling occurrence and abundance, appears as a promising solution to improve the inference of microbial associations.

The low prevalence of OTUs in metagenomic datasets greatly limits their overall analysis. In light of the results obtained, we believe that advances in the discovery of microbial associations should be made through the systematic integration of the available information in the models. The first attempts to develop statistical models incorporating prior information in the context of metagenomic analysis showed promising results [25]. From a statistical point of view, this approach can be improved by using for instance a Bayesian framework. From a a biological point of view, this approach would benefit from the development of a database dedicated to microbial interactions. Open and shared microbiota datasets, like those present in the Qiita collaborative platform (qiita.ucsd.edu), could be used to benchmark statistical models and to feed such database to improve knowledge on microbiota.

## Supporting information

**S1 Fig.**
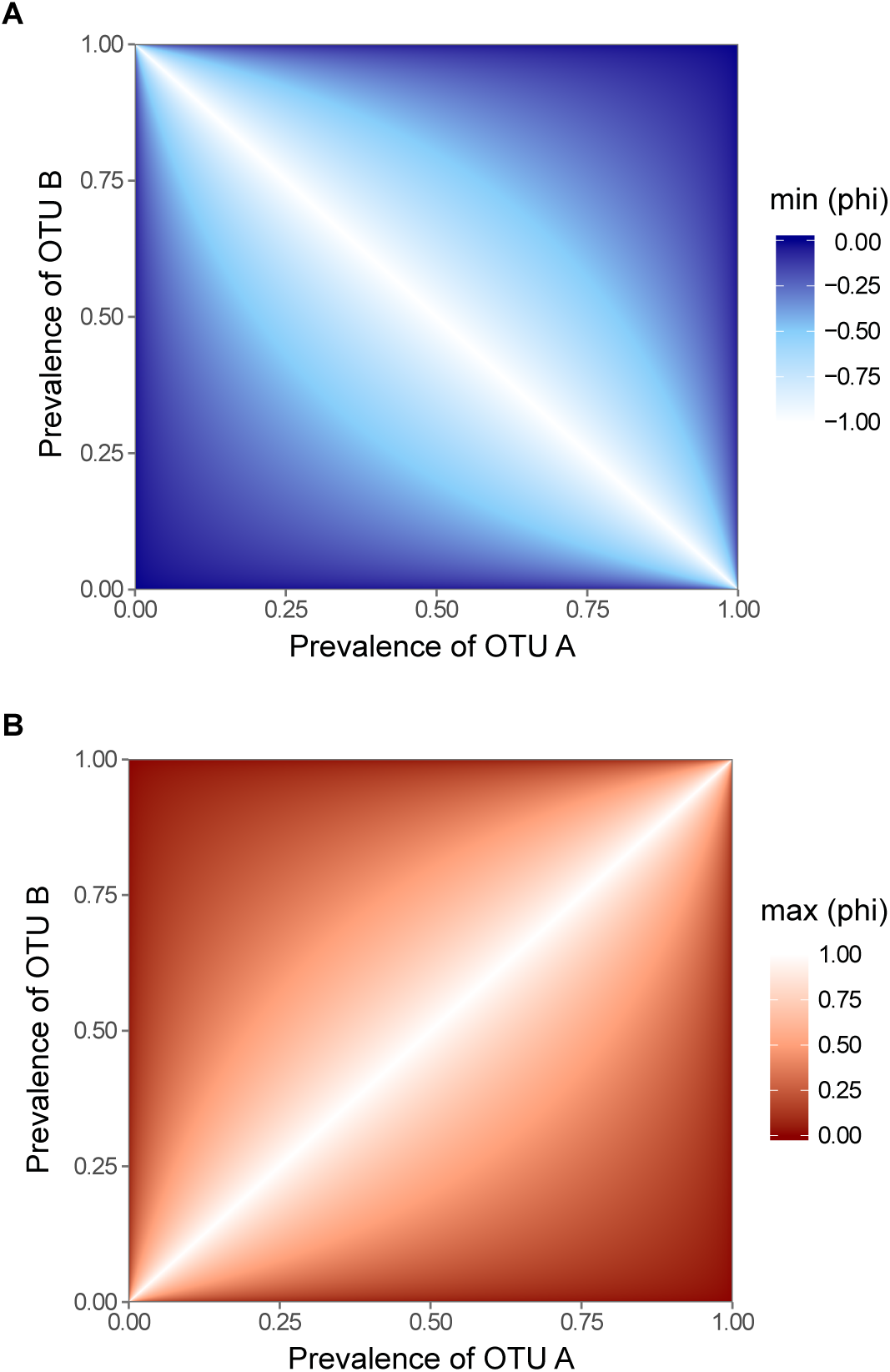
Extremum of the Phi coefficient as a function of prevalence. Minimum (**A**) and maximum (**B**) of the Phi coefficient between two OTU occurrence data as a function of their prevalence. Computed from Eq (2).

**S2 Fig.**
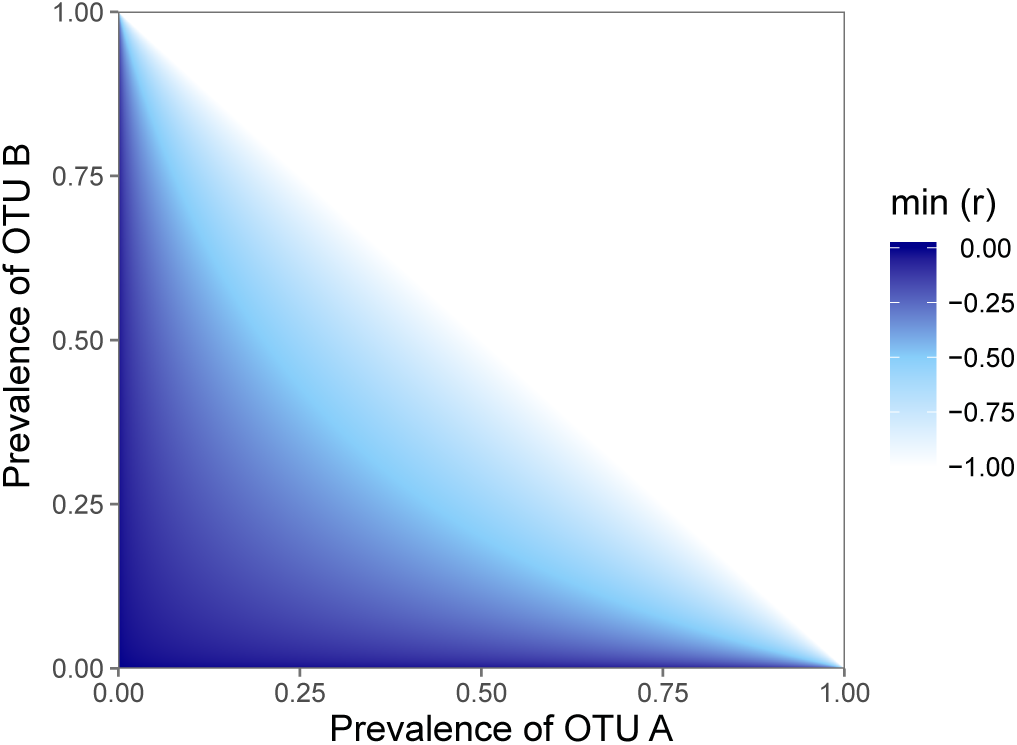
Minimum of the Pearson r correlation as a function of prevalence. Minimum of the Pearson *r* correlation coefficient between two OTU read abundance data as a function of their prevalence. Computed from Eq (5).

**S3 Fig.**
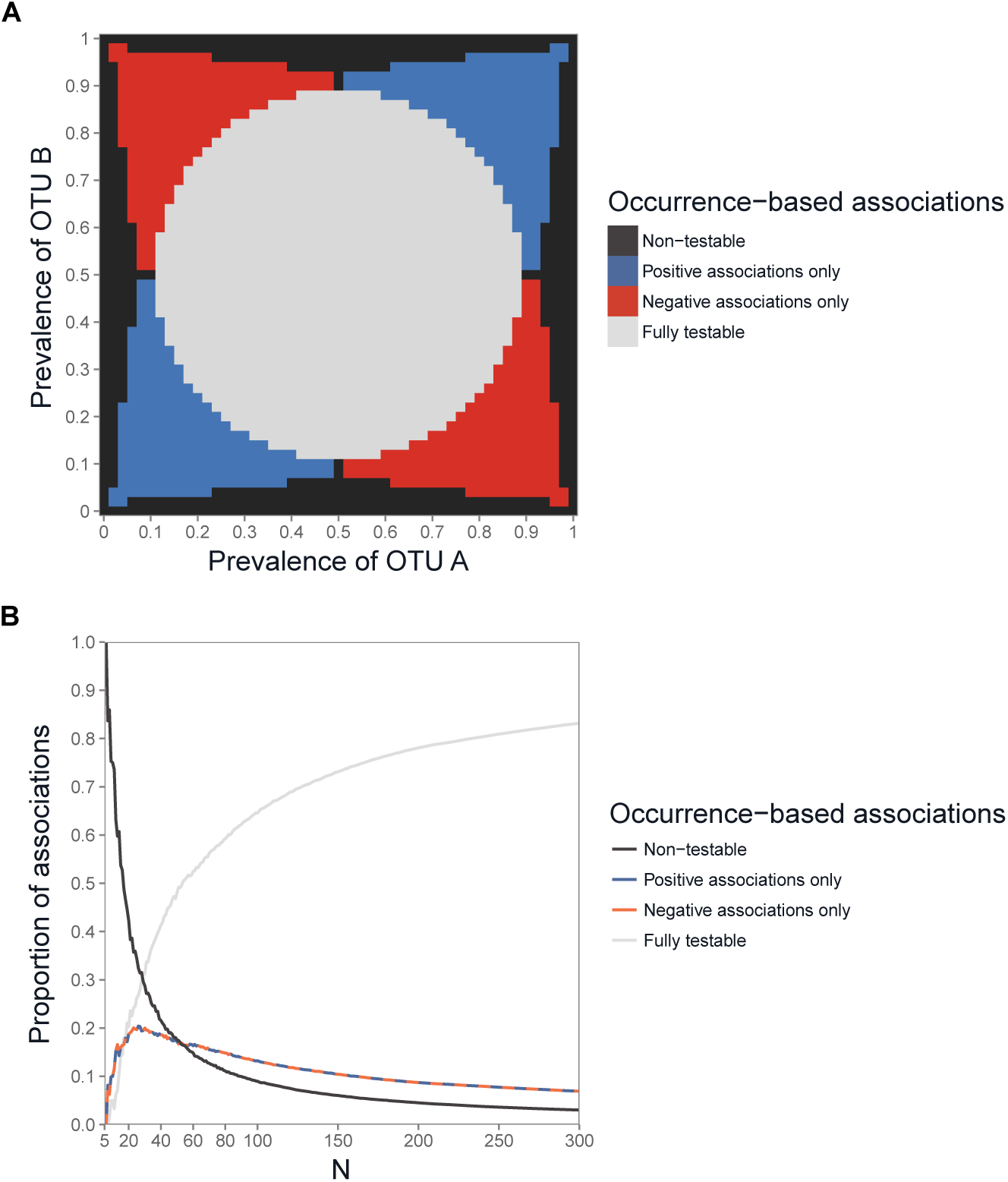
Fisher test testability of associations. Testability zones (at 5%) defined by OTU prevalence when *N* = 50 (**A**), and the proportion of testable associations as a function of *N* assuming prevalence follows a uniform distribution (**B**).

**S4 Fig.**
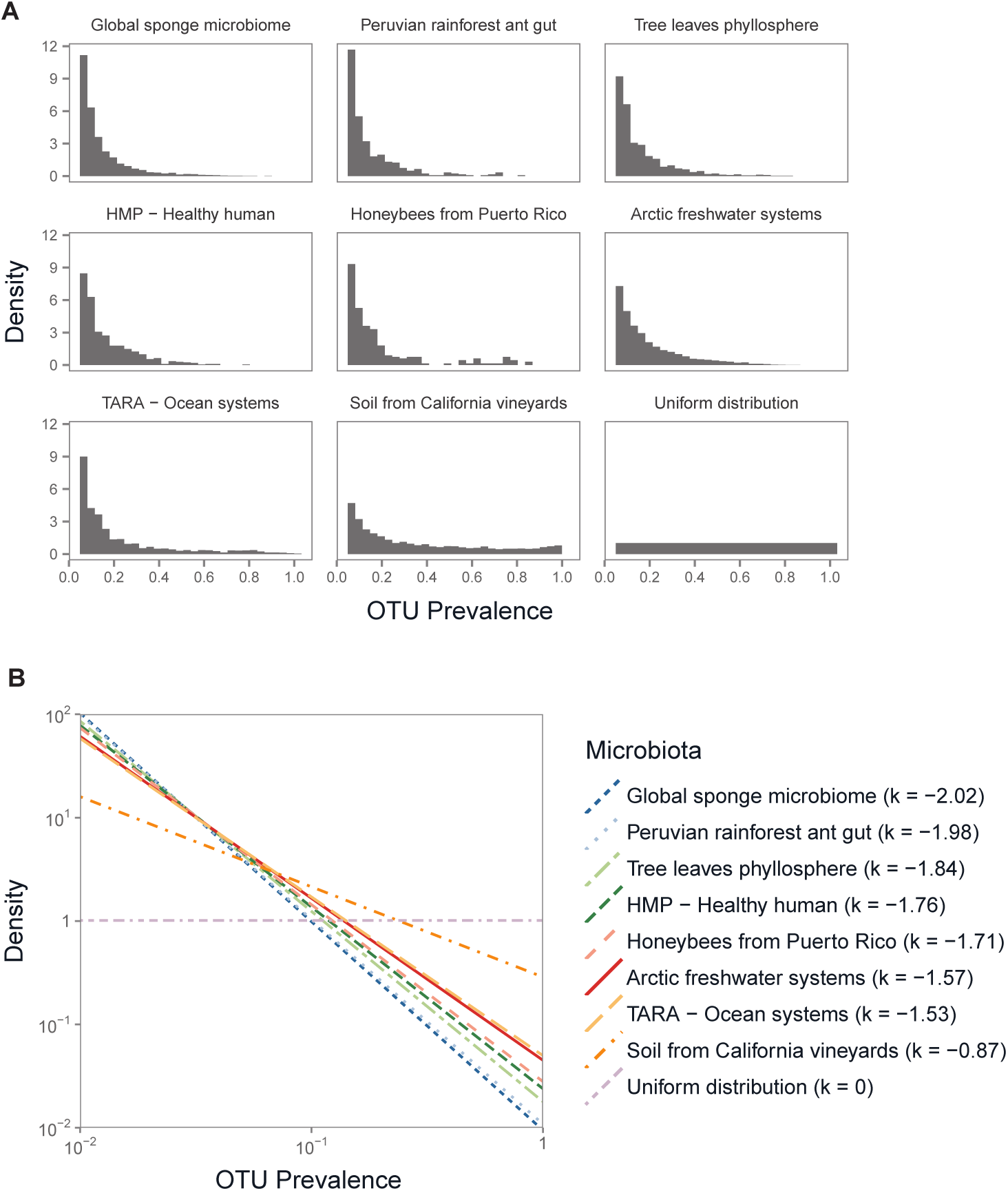
Prevalence structure of microbial community. (**A**) Histograms of OTU prevalence in several microbiota characterized by 16S rRNA sequencing. The microbiota are described in Part 5 of S1 Appendix). (**B**) Probability density function of the same prevalence data (log-log scale), which were fitted to a truncated power law distribution; the power law coefficient *k* was estimated by maximizing the log-likelihood.

**S5 Fig.**
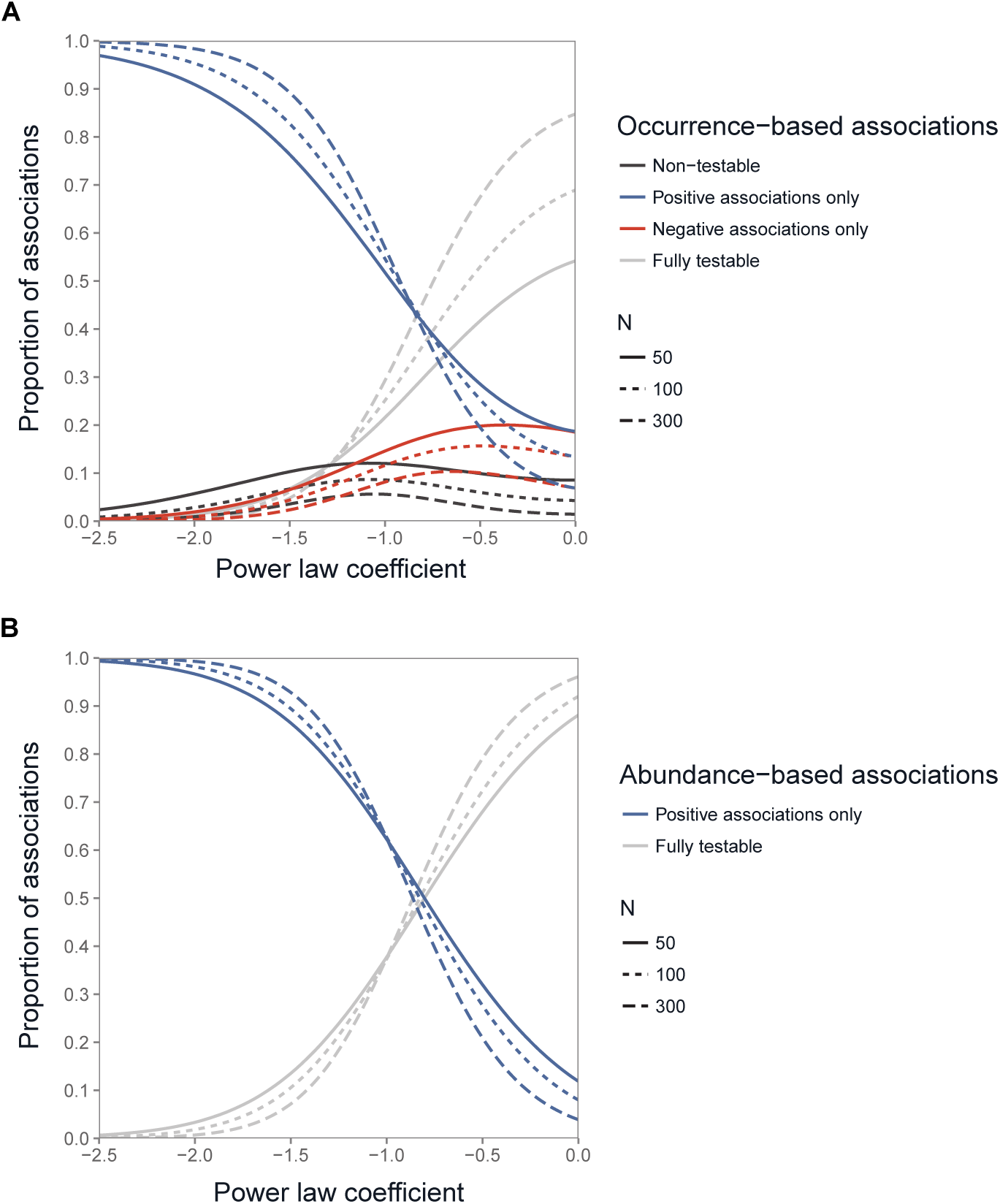
Proportion of testable associations as a function of the power law coefficient *k*. Proportion of testable associations as a function of *k* when *N* = 50, 100, or 300 for the occurrence data (**A**) and for the read abundance data (**B**).

**S1 Appendix. Supplementary Material.**

## Acknowledgments

The work was funded by two INRA metaprogrammes: Meta-omics of microbial ecosystems (MEM) and Integrated management of animal health (GISA). We thank Ioana Molnar for the mathematical advice and Jessica Pearce-Duvet for proofreading the manuscript.

## Rarity of microbial species: In search of reliable associations

Arnaud Cougoul, Xavier Bailly, Gwenaël Vourc’h, Patrick Gasqui

## S1 Appendix.

### Supplementary Material

In the supplementary material below, we describe how we established our thresholds for occurrence data (i.e., represented by binary variables) and read abundance data (i.e., represented by positive continuous variables). We then discuss the link between the two threshold types. Finally, we describe the 16S data from several microbial communities that we used to characterise prevalence patterns.

First, we present the notation and decomposition of variance and covariance as a function of OTU co-occurrence.

### 1. Notation and decomposition of variance and covariance

We consider two OTUs whose abundances are modelled by two random variables, *X*_*A*_ and *X*_*B*_ (Table 1). Our threshold is based on presence or absence of OTU, so we created a contingency table whose categories are defined by variable presence or absence.

**Table 1a.**
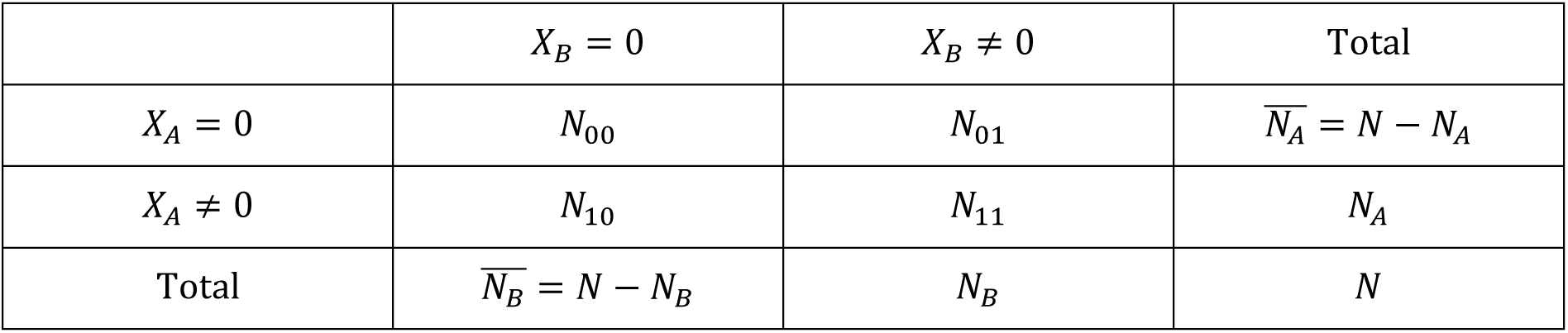
Contingency table of the presence/absence of two OTU read-abundance variables *X*_*A*_ and *X*_*B*_ where the entries are sample counts.

| | $X_B = 0$ | $X_B \neq 0$ | Total |
| --- | --- | --- | --- |
| $X_A = 0$ | $N_{00}$ | $N_{01}$ | $\overline{N_A} = N - N_A$ |
| $X_A \neq 0$ | $N_{10}$ | $N_{11}$ | $N_A$ |
| Total | $\overline{N_B} = N - N_B$ | $N_B$ | $N$ |

**Table 1b.**
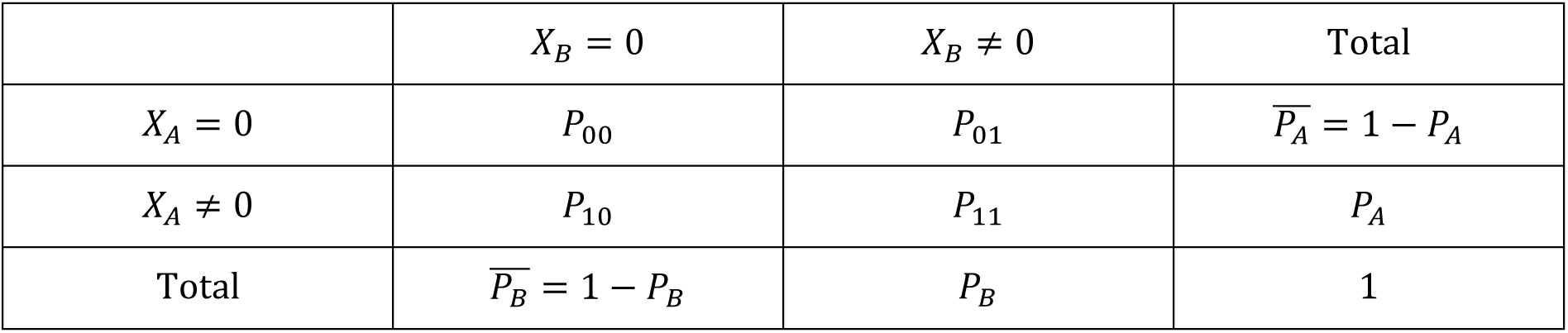
Contingency table of the presence/absence of two OTU read-abundance variables *X*_*A*_ and *X*_*B*_ where the entries are proportions.

*N* is the number of microbiota samples; *N*_00_ is the number of co-absences of *X*_*A*_ and *X_B_; N*_11_ is the number of co-occurrences of *X*_*A*_ and *X*_*B*_; and *P*_11_ = *N*_11_/*N* is the proportion of co-occurrences of the two OTUs. *P*_*A*_ = *N*_*A*_/*N* and *P*_*B*_ = *N*_*B*_/*N* are the marginal probabilities of *X*_*A*_ and *X*_*B*_, respectively (i.e., individual OTU prevalence). Since the OTUs are observed at least once, *P*_*A*_, *P*_*B*_ ∈ [1/*N*, 1].

We can calculate the mean and estimated variance of *X*_*A*_ and *X*_*B*_ using the non-zero values of *X*_*A*_ or *X*_*B*_. Consequently, 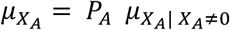, and 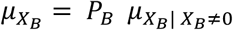.

The estimated variances of *X*_*A*_ and *X*_*B*_ can be calculated as follows:

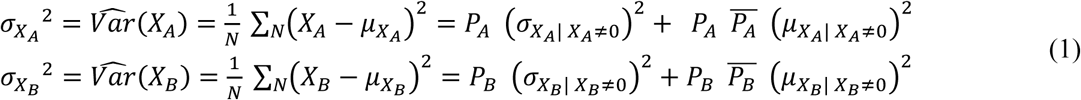

The estimated covariance of *X*_*A*_ and *X*_*B*_ can be decomposed based on whether or not *X*_*A*_ and *X*_*B*_ co-occur (i.e., *X*_*A*_ and *X*_*B*_ are non-null or not). If 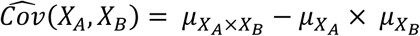, then

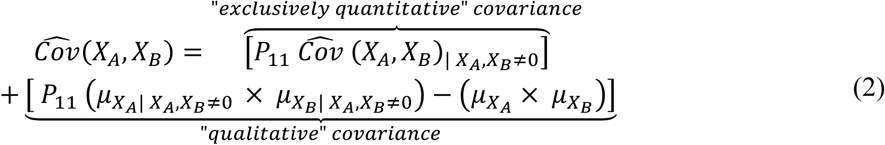

#### “Exclusively quantitative” covariance

When the data are reduced into binary variables, 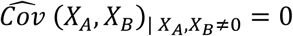 because {*X*_*A*_| *X*_*A*_, *X*_*B*_ ≠ 0} and {*X*_*B*_| *X*_*A*_, *X*_*B*_ ≠ 0} are constants. Then 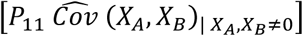 is part of the covariance of *X*_*A*_ and *X*_*B*_ only because of the quantitative aspect of data.

#### “Qualitative” covariance

The second part of the covariance 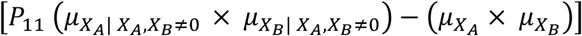 is the difference between the mean product for the whole population and the mean product for the co-occurring elements only. Consequently, it can be explained by OTU co-occurrences (qualitative in nature).

When the data are reduced into binary variables (based on equations (1) and (2)):

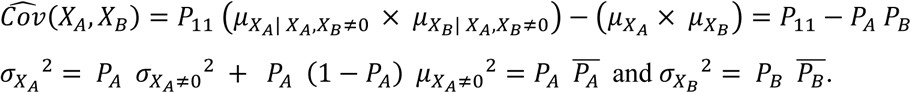

Therefore, the correlation of *X*_*A*_ and *X*_*B*_, 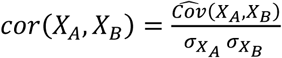, will depend only on *P*_11_, *P*_*A*_, and *P*_*B*_.

### 2. Threshold method for binary data

Our method is based on the properties of discrete statistics. As binary data are discrete data, statistical tests have discrete distributions, as do *p*-values. Moreover, the minimum observable *p*-value for fixed marginal values can be higher than the alpha level (usually set to 5%), which means the test yields useless results [1,2]. In other words, for two OTUs with fixed prevalence, if all the possible values of an association index fall within the expected confidence interval, the association is simply not testable. Below, we will illustrate how OTU prevalence can thus shape potential correlations.

In this section, we detail how we developed our threshold method for binary data (i.e., OTU occurrence). First, we describe the association index used and show that it is bounded. Second, we present how we defined its testability. Third, we examine the consequences of our threshold method for network inference. Fourth, we present the testability limits on Fisher’s exact test as a function of prevalence.

#### 2.1. Measure of associations for binary data

The combinatorics that ensue from the hypergeometric law provide only simulated solutions for determining the testability of associations. In contrast, the Phi coefficient [6] can be used to establish equations for exploring association testability and give an analytical solution. The Phi coefficient is mathematically related to the common chi-square test. Since Fisher’s exact test and Pearson’s chi-square test are asymptotically equivalent, we used the Phi coefficient as the basis for our threshold method. Moreover, we showed that the testability results were equivalent for both tests (see section 2.7 and S1 Fig 3). Phi is also equivalent to the Pearson correlation coefficient in situations with binary data (coded by 0 and 1), a property that was helpful when extending our threshold method to quantitative situations (see sections 3 and 4).

Consider two random binary variables, 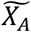 and 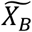, which represent the presence or absence of two OTUs. Working from Table 1, the Phi coefficient for the association between 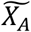 and 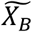 is calculated as follows:

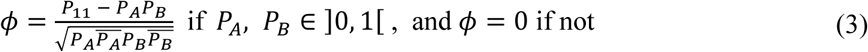

#### 2.2. Bounds of the Phi coefficient as a function of prevalence

Based on the Boole–Fréchet inequality for logical conjunction, for the marginal probabilities *P*_*A*_, *P*_*B*_ ∈]0, 1[, it follows that

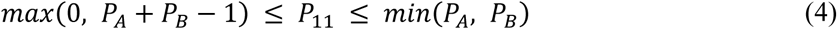

Given equations (3) and (4) and because *ϕ* is a continuous and monotonic function of *P*_11_:

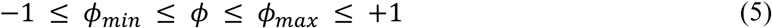

where

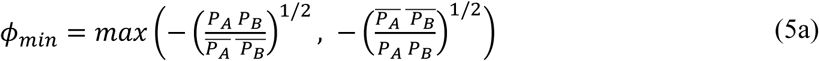

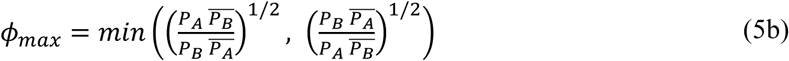

[7]

Therefore, *ϕ* is bounded and *ϕ*__min__ and *ϕ*_*max*_ depend exclusively on *P*_*A*_ and *P*_*B*_.

#### 2.3. Distribution of the Phi coefficient under the null hypothesis of independence

Under the null hypothesis (*H*_0_) that the occurrences of two OTUs, 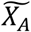 and 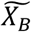, are independent, *ϕ* can be determined thanks to the Pearson’s chi-squared test: *ϕ*^2^ = *χ*^2^/*N*, where *N* is the total number of observations and *χ*^2^ is the chi-squared statistic for a 2×2 contingency table whose data follow a chi-squared distribution and for which there is 1 degree of freedom [8].

Since we know the distribution of *ϕ*, we can obtain the confidence interval at an alpha level of α. The confidence interval of a 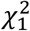 distribution is 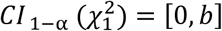, where *b* is defined by 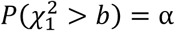 (e.g., for α = 5%, *b* ≈ 1.96^2^ ≈ 3.84).

The confidence interval of *ϕ* at an alpha level of α can be calculated as follows:

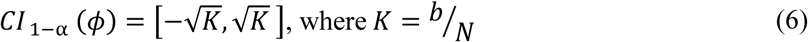

#### 2.4. Determining the testability of occurrence-based associations

We now examine the testability of the Phi coefficients calculated from pairs of OTU prevalence values. We do so by determining if the extrema of Phi occur within the confidence interval. There are two ways in which we may have trouble detecting significant associations:

A. If 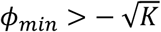, then we will not be able to detect a significant negative association.
B. If 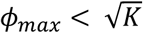, then we will not be able to detect a significant positive association.

As *ϕ*_*min*_ and *ϕ*_*max*_ depend exclusively on *P*_*A*_ and *P*_*B*_, we now consider the conditions under which *P*_*A*_ and *P*_*B*_ adopt problematic values.

We can split the first case (**A**) in two subcases because *ϕ*_*min*_ can have two different values depending on the specific values of *P*_*A*_ and *P*_*B*_:

A1) If *P*_*A*_ + *P*_*B*_ < 1, then *max*(0, *P*_*A*_ + *P*_*B*_ − 1) = 0. Based on equations (3), (4), and (5a),

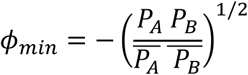
A2) If *P*_*A*_ + *P*_*B*_ ≥ 1, *max*(0, *P*_*A*_ + *P*_*B*_ − 1) = *P*_*A*_ + *P*_*B*_ − 1. Based on equations (3), (4), (5a),

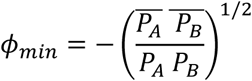

We can then resolve the inequation 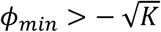.

**A1)** For 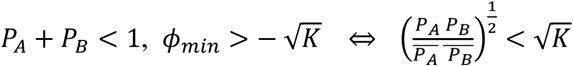

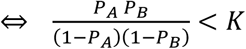 (all variables are positive)

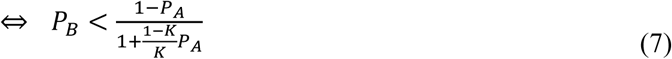

**A2)** For 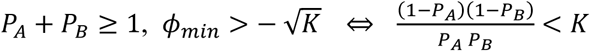

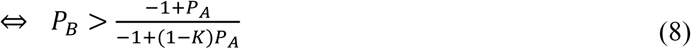

If inequations (7) or (8) are true, a negative association cannot be detected.

The second case (**B**) can be similarly split up because *ϕ*_*max*_ can also have two values:

B1) If *P*_*A*_ ≤ *P*_*B*_, then *min*(*P*_*A*_, *P*_*B*_) = *P*_*A*_. Based on equations (3), (4), and (5b),

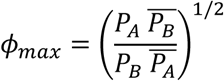
B2) If *P*_*A*_ ≥ *P*_*B*_, then *min*(*P*_*A*_, *P*_*B*_) = *P*_*B*_. Based on equations (3), (4), and (5b),

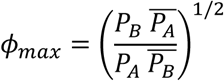

We can now solve the inequation 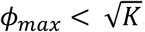.

**B1)** If 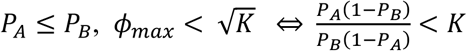

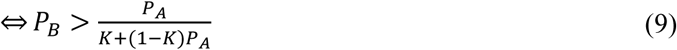

**B2)** If 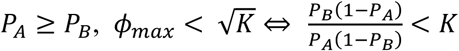

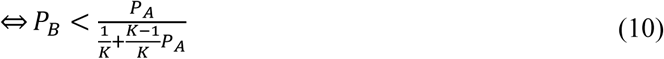

If inequations (9) or (10) are true, a positive association cannot be detected.

Using the four inequations (7), (8), (9), and (10), we can delimit zones within which there is full, partial, or no testability. The characteristics of the tests in these zones will be detailed in the introduction to the next section.

For the two OTUs, *P*_*A*_ and *P*_*B*_ form a [1/*N*, 1]^2^ square (Figure 1 below); 1/*N* is the smallest observable value. The testability zones in this square can be defined using four border functions that result from the inequations:

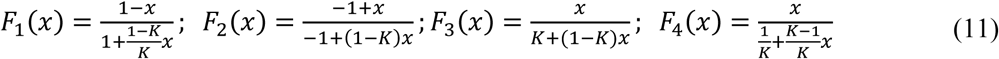

**Figure 1.**
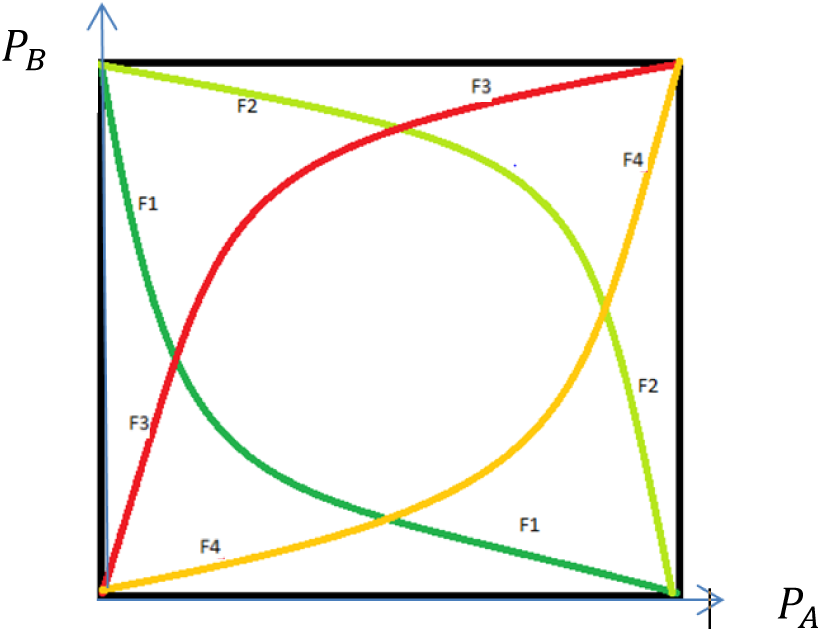
The four border functions delimiting testability

Emerging from these border functions are four graph intersections that are defined by:

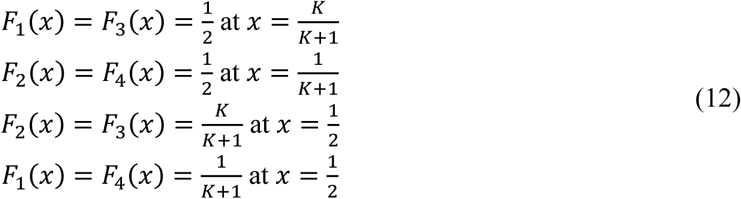

#### 2.5. Proportion of associations in each testability zone

The zones defined by the border functions (11) contain different proportions of associations that can be categorised as fully testable, partially testable, or non-testable using our threshold method. The first zone, *A*_*bilateral*_, contains associations for which both positive and negative correlations can be reliably tested. The second zone, *A*_*unilateral*_, contains associations for which only positive correlations can be reliably tested (subzone *A*_*positive*_) and for which only negative correlations can be reliably tested (subzone *A*_*negative*_). Finally, the third zone, *A*_*irrelevant*_, contains associations that cannot be reliably tested at all.

The distribution of prevalence values is treated as identical for all OTUs. Therefore, *P*_*A*_ and *P*_*B*_ have the same distribution and play symmetrical roles. However, these distribution patterns are not necessarily uniform. We examined two types of distributions—the uniform distribution and the truncated power law distribution; the latter fit the prevalence patterns of OTUs in real microbiota (see section 5).

For the uniform distribution of prevalence, the probability density function is

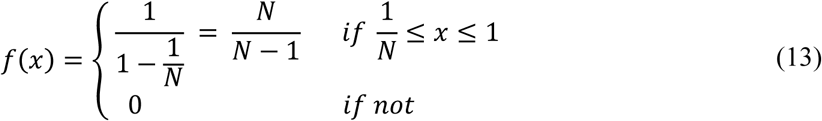

For the truncated power law distribution of prevalence, the probability density function is

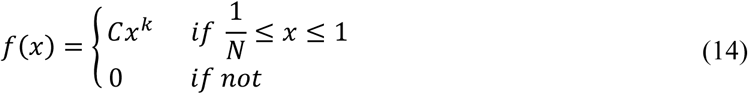

and, following normalization, we arrive at 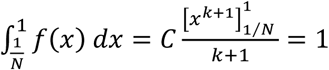, so 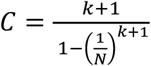.

When *k* = 0, we have a uniform distribution with the interval 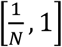.

To computationally define the different zones, analytical formulas can be used in the case of the uniform distribution but not in the case of the power law distribution. Consequently, in the latter situation, we chose to proceed by numerical integration. Since the current form of the R function *integrate* (in the stats package) does not deal well with the power law, we used a Monte Carlo approach. This consisted of generating random prevalence values in accordance with the observed prevalence distribution (see section 2.6) and counting how many fell within each of the zones.

To simplify the zone-defining equations below, we have used the following notation:

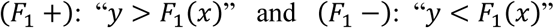

and the same notation applies in the cases of *F*_2_, *F*_3_ and *F*_4_.

∧ denotes the logical conjunctions.

From the four inequations (7, 8, 9, 10) and the border function (11), the proportions of associations that fall within each zone are determined as follows:

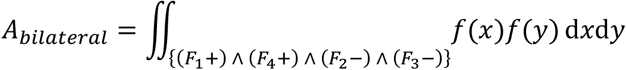

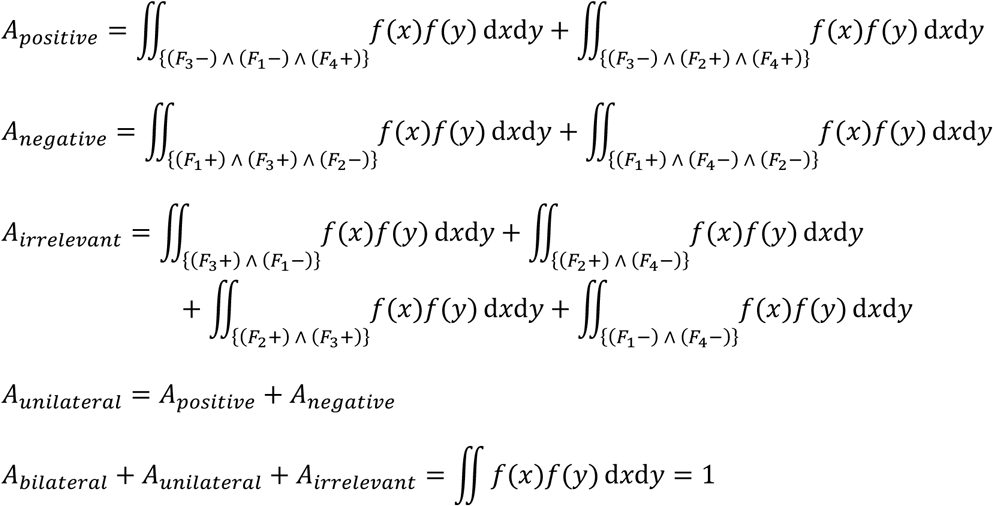

#### 2.6. Defining testability zones using a Monte Carlo method

To compute Monte Carlo integrations, it is necessary to generate random prevalence values using the observed distribution of prevalence. For the uniform distribution, many pseudorandom number generators exist. However, for the truncated power law distribution, we had to employ an inverse transformation method that is rooted in the following property:

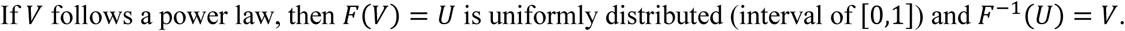

We therefore needed to define the inverse cumulative distribution function. Let *F* be the cumulative distribution function of the truncated power law distribution as defined in (14).

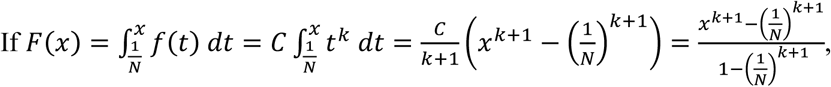

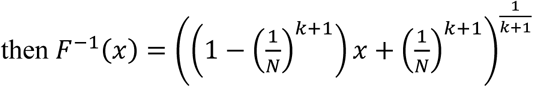

We can then generate a power law distribution from a uniform distribution using the following equation:

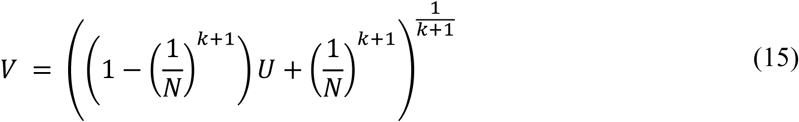

#### 2.7. Testability limits on Fisher’s exact test

Co-occurrence networks are commonly reconstructed using the hypergeometric law that underlies Fisher’s exact test [3–5].

From an observed 2×2 contingency table (Table 1), Fisher showed that the probability *P* of obtaining such a set was given by the hypergeometric distribution:

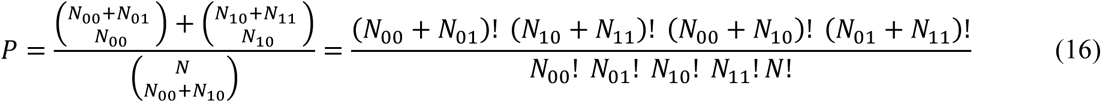

where 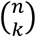 is the binomial coefficient and ! indicates the factorial.

This equation can be written according to *N*_*A*_, *N*_*B*_, *N* and *N*_11_:

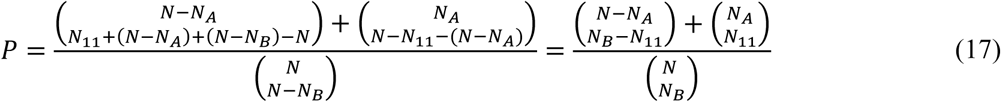

Based on the Boole–Fréchet inequality for logical conjunction, for the marginal counts *N*_*A*_, *N*_*B*_ ∈]0, *N*[, it follows that

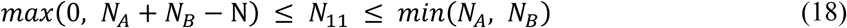

We have two extreme situations:

a. Observe the minimum number of co-occurrences, *N*_11_ = *min*(*N*_11_) = *max*(0, *N*_*A*_ + *N*_*B*_ − N)
b. Observe the maximum number of co-occurrences, *N*_11_ = *max*(*N*_11_) = *min*(*N*_*A*_, *N*_*B*_)

We can calculate the probability *P* associated with these two situations **a)** and **b)**. A bilateral test can also be performed. As in the `fisher.test` function of R, the p-value is computed by summing the probability for all table with probabilities less than or equal to that of the observed table.

For two given OTUs with prevalence 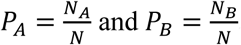 and 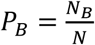, we have 4 possibilities in the testability limits on Fisher’s exact test:

- If the p-values associated with the two configurations **a)** and **b)** are lower than the alpha level (5%), the two extremes situations **a)** and **b)** correspond to significant associations. We have no limit on the test.
- If the p-value associated with the configuration **a)** is greater than the alpha level, then we will not be able to detect a significant negative association.
- If the p-value associated with the configuration **b)** is greater than the alpha level, then we will not be able to detect a significant positive association.
- If the p-values associated with the configurations **a)** and **b)** are greater than the alpha level, then we will not be able to detect a significant positive or negative association.

### 3. Threshold method for quantitative data

In this section, we detail how we developed our threshold method for quantitative data (i.e., OTU read abundance). First, we introduce our system of notation and the primary elements of our proof. Second, we present the situation, in which correlations are bounded by an excess of zeroes, and describe the minimum correlation value. Third, we show how we defined association testability. Finally, we examine the consequences of our threshold method for network inference.

#### 3.1. Introduction

In this section, *X*_*A*_ and *X*_*B*_ are two random variables that represent quantitative data. The Pearson correlation coefficient [9] is used to characterise the pairwise associations in OTU read abundance. We were specifically interested in understanding how the number of zeroes in the data could influence the correlation coefficient.

We use same notations as in section 1. 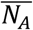 and 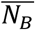 represent the number of zeros associated with *X*_*A*_ and *X*_*B*_, respectively. *N*_00_ is the number of co-absences of *X*_*A*_ and *X*_*B*_, and *N*_11_ is the number of co-occurrences.

Based on Table 1 and the Boole–Fréchet inequalities, we can deduce the following:

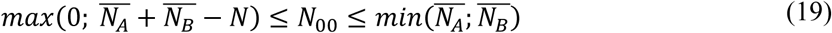

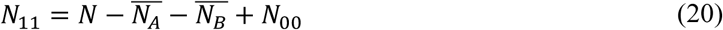

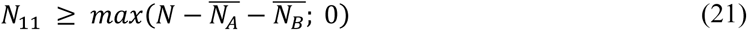

For pairs of 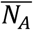 and 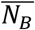 we distinguish two cases:

i. 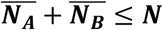 The number of zeros is sufficiently low such that there are no raw restrictions on possible correlations. Indeed, it is simple to build two non-restricted correlations that approach infimum −1 and supremum +1:

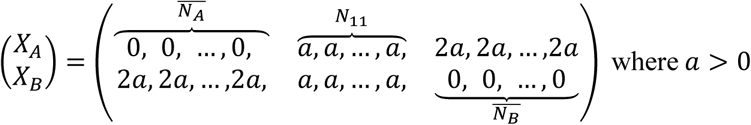 In this case, the correlation coefficient is *r* = −1.

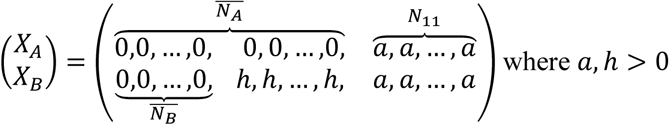 The correlation tends toward the supremum, 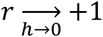 or 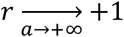.
ii. 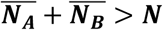 Based on equations (19) and (21), 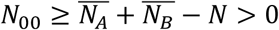 and *N*_11_ ≥ 0. Consequently, *N*_11_ can equal zero, meaning that there are enough zeros associated with *X*_*A*_ and *X*_*B*_ that *X*_*A*_ and *X*_*B*_ may not co-occur. In this situation, information on quantitative correlations is degraded. We can prove that *r*, the Pearson correlation coefficient, has a minimum, *r*_*min*_, that is different from −1:

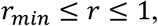

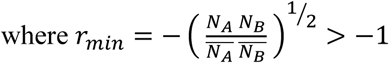

#### 3.2. Determining the lower bound of the Pearson correlation coefficient

Given 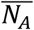 and 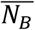 we wished to determine the minimum possible correlation between *X*_*A*_ and *X*_*B*_. We highlight that a lower bound of the Pearson correlation exists between two positive variables and prove that it can be reached under certain conditions.

For the association between *X*_*A*_ and *X*_*B*_, the Pearson correlation coefficient is calculated as follows:

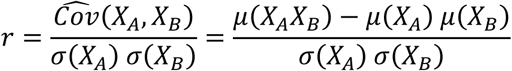

If *X*_*A*_, *X*_*B*_ ≥ 0, then *μ*(*X*_*A*_*X*_*B*_) ≥ 0 and

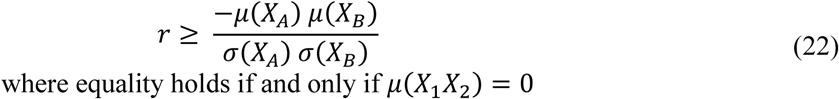

Consequently, the mean of *X*_*A*_*X*_*B*_ is null if and only if there are no co-occurrences. In other words,

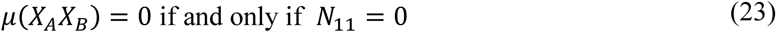

- If *μ*(*X*_1_*X*_2_) = 0, then ∑ *X*_1_*X*_2_ = 0. Each element of the sum are positive then ∑ *X*_1_*X*_2_ = 0 imply that all elements are null and there are no co-occurrences (i.e., *N*_11_ = 0).
- If there are no co-occurrences, then *X*_1_*X*_2_ = 0 and *μ*(*X*_1_*X*_2_) = 0.

From equations (22) and (23), we can conclude that

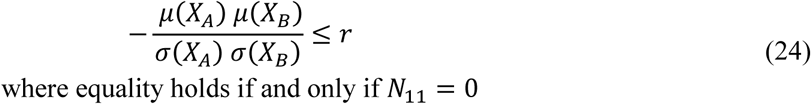

Moreover, if 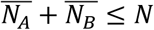, then, from equation (21), we know that *N*_11_ ≠ 0. Therefore,

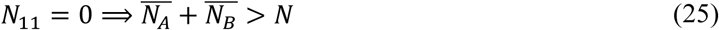

We now want to control 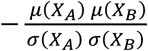 and find its minimum. We therefore maximise 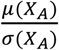 and 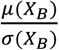 separately. *μ*/*σ* corresponds to the inverse coefficient of variation.

#### 3.3. Maximising the inverse coefficient of variation

Below, we illustrate how to maximise the inverse coefficient of variation for *X*_*A*_. We will show that 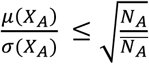.

We can express variance using the König–Huygens formula:

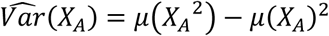

If *μ*(*X*_*A*_) ≠ 0, then

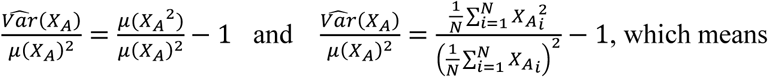

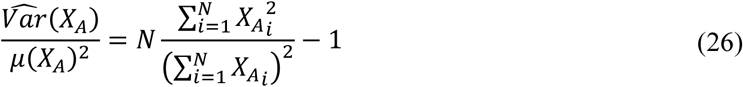

We are now interested in 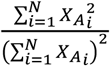, and we will show that 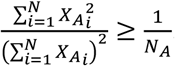.

Let *V, W* be two vectors of ℝ^*N*^. As per the Cauchy–Schwarz inequality,

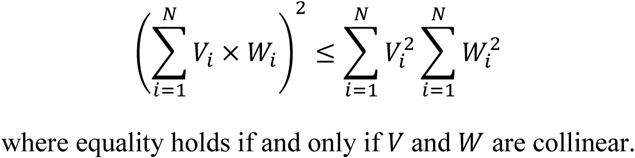

Let *V* = *Y* be the vector of non-null elements of *X*_*A*_ (for *Y*, vector size is equal to *N*_*A*_); *W* = 1_*N*_*A*__, a constant vector of size *N*_*A*_. In this case, the Cauchy–Schwarz inequality becomes the following:

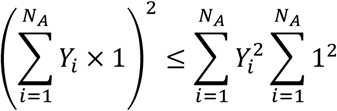

where equality holds if and only if *Y* = *λ* × 1_*N*_*A*__, where *λ* > 0 (i.e., *Y* is a constant vector).

As 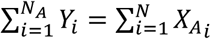 and 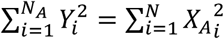, 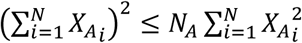, then

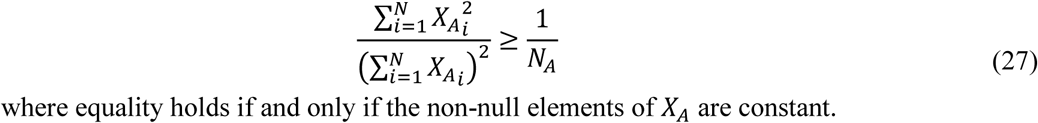

Based on equations (26) and (27), we now observe that

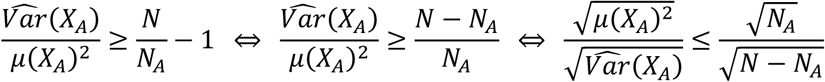

Finally,

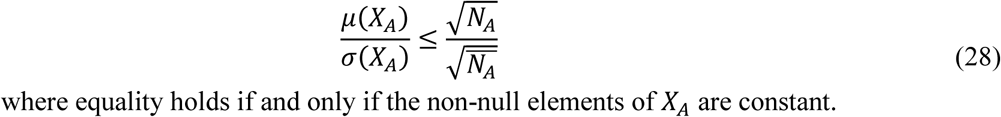

The maximum occurs where 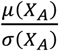 is 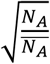.

The approach is equivalent for *X*_*B*_, so we can conclude that

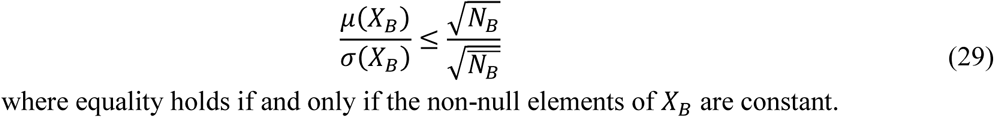

#### 3.4. Determining the minimum Pearson correlation coefficient when there are many zeros

Based on equations (24), (28), and (29),

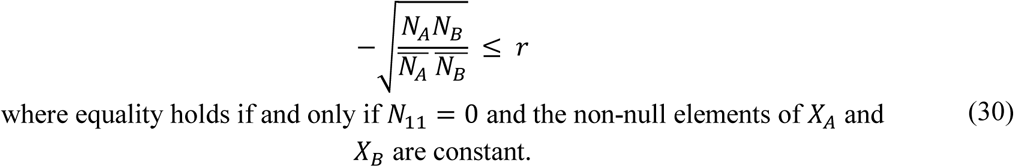

It therefore stands to reason that

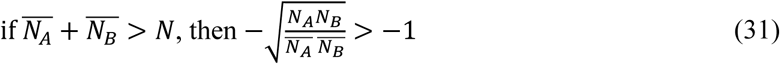

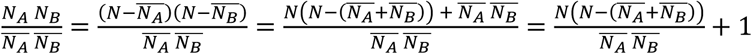

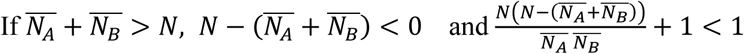

Therefore, 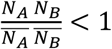 and 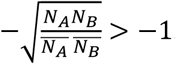.

Finally, based on equations (30) and (31), when 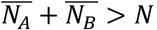,

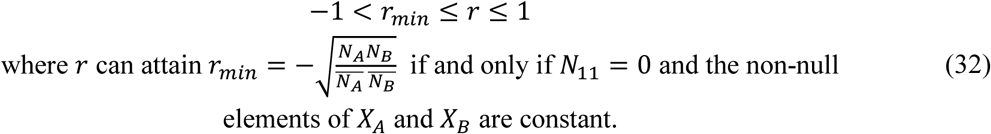

#### 3.5. Constraints on the testability of the Pearson correlation coefficient

When *X*_*A*_ and *X*_*B*_ follow two uncorrelated normal distributions, 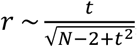, where *t* is a Student’s t statistic with degrees of freedom *N* − 2. We can then determine a confidence interval: 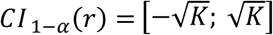, where *K* depends on *α* and *N*.

Returning to our measures of OTU prevalence, if 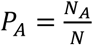 and 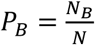, then 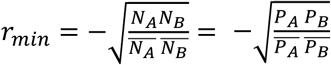. The constraint is the same as in the case of binary data.

If *r*_*min*_ falls within the confidence interval, we can conclude that negative associations cannot be detected.

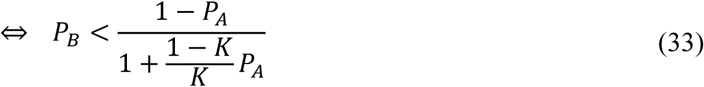

Accordingly, if inequation (33) is true, then negative associations are not testable.

The border function that defines the testability zones in the square formed by *P*_*A*_ × *P*_*B*_ is as follows:

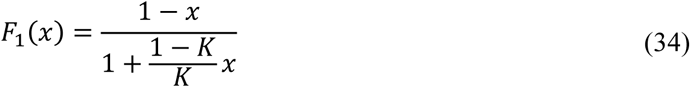

#### 3.6. Proportion of associations in each testability zone

Using the border function (34), we observed that two zones existed. The first zone, *A*_*bilateral*_, contains associations for which both positive and negative correlations can be reliably tested. The second zone, *A*_*positive*_, contains associations for which only positive correlations can be reliably tested. As for the binary data (sections 2.5 and 2.6), we explored the testability of abundance-based associations using the uniform distribution and the truncated power law distribution. In the latter case, we again employed a Monte Carlo approach.

Based on the border function (34), the proportions of associations that fall within each zone can be determined as follows:

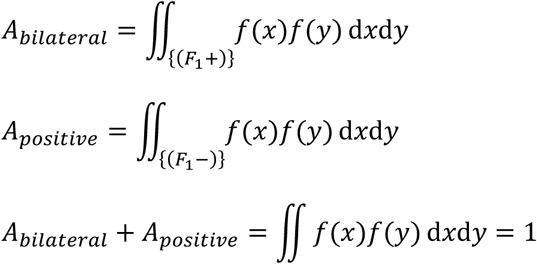

(Same notation as in section 2.5)

### 4. Similarity of the Phi and Pearson correlation coefficients

In this section, we show that testability constraints tend to be similar with both occurrence and abundance data. We also examine the degree of correlation between the correlation coefficients calculated using the two data types.

#### 4.1. Testability constraints on occurrence and abundance data

The distribution of the correlation coefficient for two normally distributed independent variables is 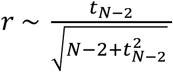.

As 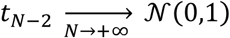 (i.e., there is distribution convergence) and 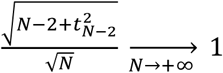, then 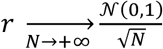. Since the distribution of the square of the Phi coefficient is 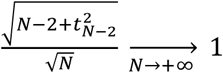 under the null hypothesis of independence, the Pearson correlation coefficient will asymptotically attain the same confidence interval as the Phi coefficient: their lower bounds converge upon 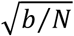 (sections 2.3 and 3.5).

We now underscore that the Phi and Pearson correlation coefficients have the same lower bound when the two OTUs have low levels of prevalence: 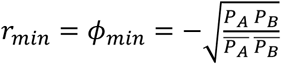

When *N* is large enough, the testability of positive associations will be the same for binary data and quantitative data. This pattern will be all the more pronounced given that, in real microbiota, OTU prevalence is greatly skewed to the right: positive associations represent the majority of associations to be tested.

#### 4.2. Correlation between Phi and Pearson coefficients

In section 1, we showed that variance can be decomposed in a quantitative part and a qualitative part (equation (2)). Here, we use the results of a simulation to explore how the strength of the correlation between the values of the Phi coefficient and the Pearson coefficient is related to OTU prevalence. We are most interested in what happens when prevalence is low.

OTU abundances *X*_*A*_ and *X*_*B*_ are modelled by a zero-inflated Poisson (ZIP) distribution using the following probability mass function:

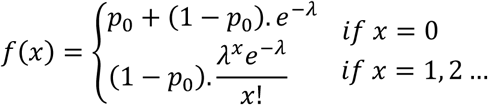

where the probability of structural zeros, *p*_0_, is the result of a Bernoulli process and *λ* is the mean of the Poisson portion of the distribution (i.e., the Poisson parameter). In the simulation, *X*_*A*_ and *X*_*B*_ had the same values for *p*_0_ and *λ*.

The probability of structural zeros *p*_0_ represents the complementary probability of prevalence *P*, i.e. *p*_0_ = 1 − *P*. As *p*_0_ increases (i.e., prevalence decreases), the correlation between the Phi coefficient and the Pearson coefficient increases (Figure 3A in the article). The correlation also strengthens as *λ* increases. When prevalence is below 0.25, the correlation is greater than 0.75 for all values of *λ*.

If OTU prevalence follows a ZIP distribution, we can conclude that the values of the Phi coefficient and the Pearson coefficient will be correlated, especially when OTU prevalence is low.

### 5. Distribution of OTU prevalence in real microbiota

To characterise actual OTU distribution patterns, we employed data from the QIITA database (qiita.ucsd.edu) and the TARA Ocean Project (ocean-microbiome.embl.de) [10]. The biom files were processed using the R package *biomformat*. We deliberately chose different kinds of microbiota so as to represent as wide a diversity of microbial communities as possible (Table 2). We used OTU rather than species tables.

**Table 2.**
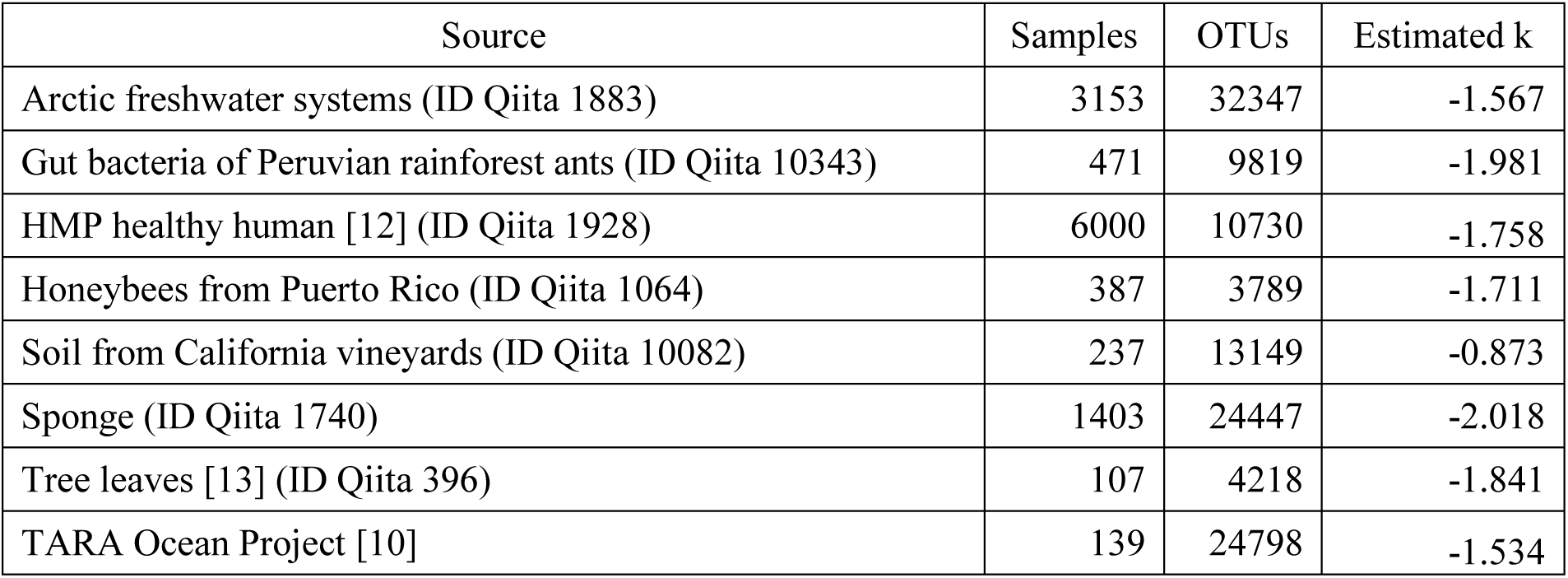
Sources of the microbiota we analysed and the associated number of samples, number of OTUs, and estimates of the power law coefficient k.

The prevalence values were fitted to a truncated power law distribution as described by equation (14), and the power law coefficient k was estimated by maximizing the log-likelihood [11].

